# A sneak peek at the high altitude adaptation of the Ladakh populations

**DOI:** 10.1101/2024.03.31.587460

**Authors:** Urgyan Chorol, Bhagyashree Choudhury, Tsering Norboo, Nony P. Wangchuk, Gyaneshwer Chaubey, Chandana Basu Mallick

## Abstract

The physiological response to high-altitude stress (for individuals living at 2,400m and above sea level) has been evident. However, recent advances in genomics have allowed us to explore the molecular genetic basis of these adaptive responses and their relationship with physiological responses. In the current study, we focused on thirteen biological parameters to understand the adaptive response to the high altitude of populations living across regions of Sakti, Korzok, Hanle, Aryan and Zanskar valleys of Ladakh and to understand the variation between and across individuals. Interestingly, we found a negative correlation between haemoglobin levels and partial oxygen pressure, thereby testifying to higher haemoglobin levels as an adaptive response in these individuals. The lipid profiles, including cholesterol, triglyceride, LDL and HDL levels, varied significantly across the five regions studied. Notably, individuals from Sakti Valley showed higher LDL (>140) and cholesterol values. These variations in health parameters and lipid profiles may be attributed to diet and altitude adaptations specific to the region. Overall, the study provides valuable insights into the adaptations of highlanders living at different altitudes in the Ladakh region. The findings can have implications for better understanding the molecular mechanism of physiological responses of humans to high-altitude environments.

## INTRODUCTION

The hypoxic condition of high altitude imposes severe physiological responses. Studying the high altitude populations can help us better understand the genetic basis of adaptive physiological traits (Storz and Cheviron, 2021). The next-generation sequencing technologies have provided us with opportunities to unleash it. Previous studies on different high-altitude populations such as Andeans, Tibetans, Ethiopians and Sherpas have revealed that they all have different stories to tell, leading to population differences in haematological response to high-altitude hypoxia (Beall, 2014, 2006; Beall and Strohl, 2021; Scheinfeldt et al., 2012). However, though different genes are responsible for adaptation in different populations, attesting to convergent evolution (Lee and Coop, 2019; Losos, 2011), what has been shared is that most of the genes have undergone positive selection to the same selective pressure and converge on similar physiological pathways. Hence, studying genes involved in high-altitude adaptation further attests that most complex traits are polygenic and exhibit pleiotropy (Zheng et al., 2023). Among the high-altitude populations studied, the Himalayan region has been less extensively studied, restricted to only a handful of studies. Also, Asia has considerably the largest variation in terms of populations living in high altitude ranging from low to high and hence studying them can help in identifying unique physiological mechanisms crucial to high altitude adaptation.

Ladakh has a wide range of altitudes (2600-4900m), is primarily an arid landscape, and is subject to many environmental challenges. It is in the northernmost part of India. It is comprised of Leh and Kargil districts, occupied by different ethnic groups followed by Changpa, Balti, Bodh and Brokpa. The Changpa tribes are located towards the eastern part of Ladakh, an extension of the Tibetan plateau; they are also called nomadic people or pastoralists. In Ladakh, the pastoral communities settled on the extension of the Tibetan plateau were believed to have originated from ancient tribes on the Tibetan plateau (Ganjoo and Ota, 2012; Leipe et al., 2014). They move from one place to another in search of pasture and livestock. They are exposed to very harsh environmental conditions because of living at an altitude of 4500m above sea level. Multiple studies revealed that continuous exposure to harsh environments allows the population to undergo physiological and genetic (Bigham et al., 2010; Brutsaert, 2007; Droma et al., 1991; Gassmann and Muckenthaler, 2015; Huerta-Sánchez et al., 2014; Jeong and Di Rienzo, 2014; Moore, 2017; Simonson et al., 2012; Zhuang et al., 1993). Most people in the upper Changthang area are pastoralists, while populations in the lower Changthang area practice farming. Balti and Bodh live at 3000m above sea level in central Ladakh. Their occupation is also farming. The Brokpa have inhabited the western part of Ladakh along the bank of the Indus River. They are an Indo-European-speaking tribe in Ladakh, claiming to originate from Gilgit Pakistan, where the Dardic speakers are present. They practice farming and horticulture; their apricots are known for being the sweetest apricots in Ladakh. In the current study, we have studied thirteen health parameters (including anthropometric, hematological and lipid profiles) to understand the adaptive response to the high altitude of populations living across regions of Sakti, Korzok, Hanle, Aryan and Zanskar valleys. We aim to understand the within and across region diversity and how they influence their adaptive responses. In particular, we have studied their haematological, lipid, blood pressure and anthropometric profiles.

## 3. METHODOLOGY

### STUDY SELECTION

#### Study settings and participants or study selection

Our study includes a total of 458 participants. These samples were collected from five geographic regions - the Zanskar region (N=225), Aryan valley (N=85), Korzok region (N=16), Hanle region (N=72) and Sakti valley (N=60) [(Figure 1)]. Written informed consent was procured from all volunteers. Ethical approval was obtained from the District Ethical Committee Ladakh for this study and fieldwork. We have collected demographic, anthropometric and lifestyle information, including gender, age, place of residence, marital status, educational level, employment status, ethnic group, weight, height, body mass index (BMI), Partial oxygen pressure (Spo2) by pulse oximeter, heart rate, waist circumference, systolic blood pressure and diastolic blood pressure. We also collected haematological and lipid profile-related parameters, including haemoglobin, blood sugar (BS), Cholesterol, triglycerides, low-density lipoprotein (LDL), high-density lipoprotein (HDL), creatinine (Cr) etc.

**FIGURE 1:**
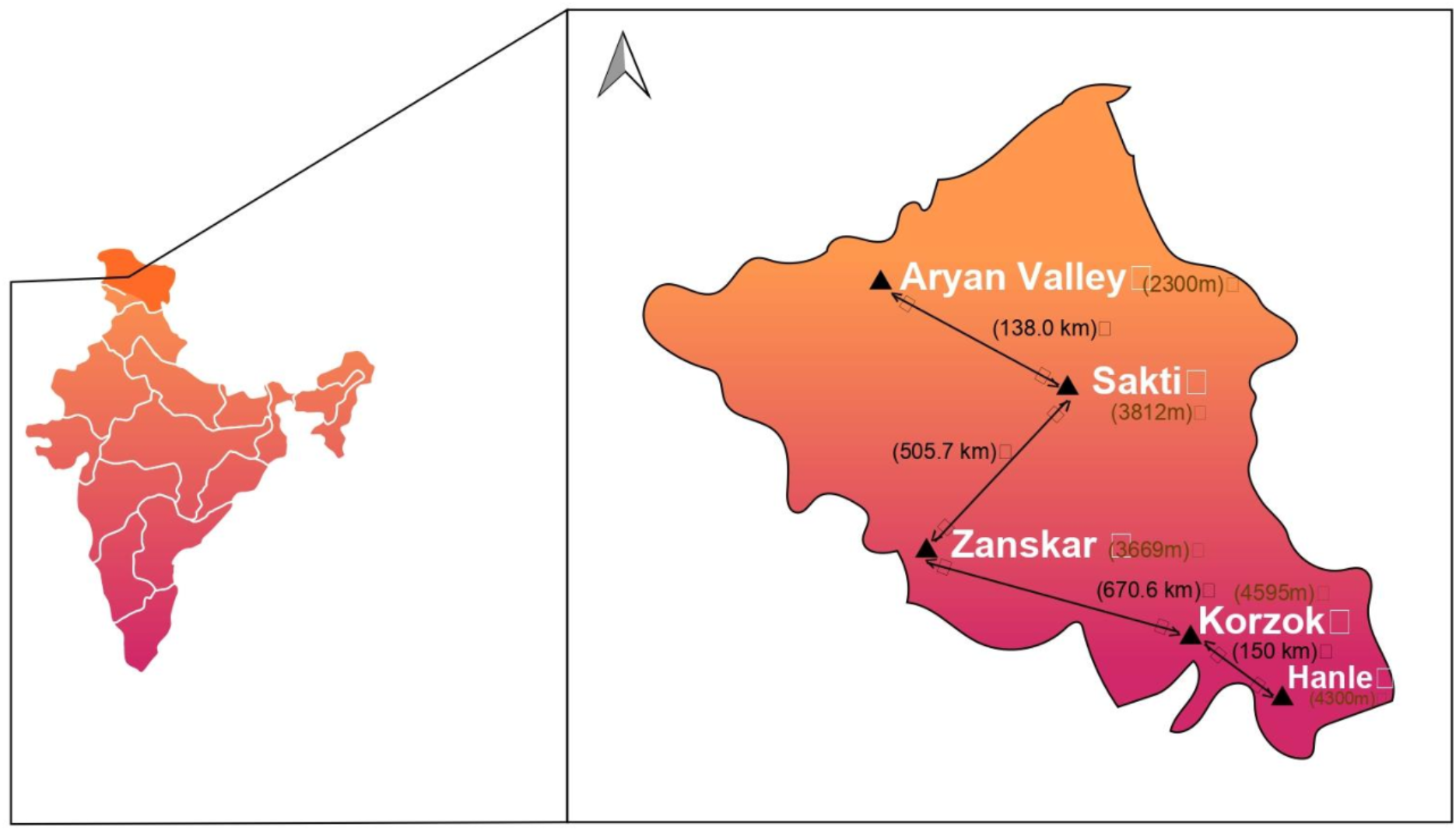
Map of Ladakh Region showing the sampling sites together with their elevations in the subdivisions.

### STATISTICAL ANALYSIS

All the analyses were carried out on GraphPad Prism version 9.5.1. Quantitative data analysis was expressed as means and standard deviations (mean±SD). One sample t-test and Wilcoxon test were used to examine whether the mean of a population is statistically different from a known or hypothesised value. One-way ANOVA with the Bonferroni significant difference test was performed to assess the significance of differences between the groups (multiple comparisons) and reduce the probability of making Type I errors. The correlation matrix tool was used to determine whether the variables were significantly, poorly or not connected. The considered statistically significant level was p<0.05.

## 4. RESULTS

### Evaluation of the anthropometric profile

When considering body composition metrics, Sakti emerges as the dominant group, exhibiting the highest values across all three parameters: body weight (59.78±9.485 kg), height (173.79±17.49 cm), body mass index (25.97±3.806 kg/m^2^) and waist circumference (83.23±9.339 cm). In the context of males, Zanskar exhibits elevated values for height (173.37±12.22 cm) and body mass index (23.88±4.91 kg/m^2^), while Sakti claims the lead for body weight (61.76±10.91 kg). Conversely, among females, Sakti showcases higher measurements across all three aspects: body weight (59±8.87 kg), height (161.48±0.068 cm), and body mass index (26.88±3.65 kg/m^2^). Among males, Zanskar records the highest mean waist circumference (84.14±11.37 cm), whereas, among females, Sakti showcases the highest value (83.91±9.652 cm) (Table 1,2 & 3).

Examining the correlation matrix, we find a subtle negative association between Korzok and Aryan regarding height (r=-0.2914), whereas weight exhibits no significant correlation. Notably, body mass index demonstrates a positive association (r=0.3415) (Table 4) between Zanskar and Korzok. When we did a one-way analysis, we detected noteworthy distinctions between Hanle and Sakti (p=0.0052)) (Table 5b). The study highlights variations in waist circumference measurements across different regions and gender. Sakti consistently displays higher measurements, while Hanle shows lower values. However, the correlation analysis does not yield any significant association for waist circumference.

A one-way ANOVA analysis was performed [Figure 2 (a, b, c and d)], and height showed non-significant differences among groups. For body weight, significant associations surface between Aryan and Korzok. The body mass index varies significantly between Zanskar and Sakti (p=0.0049) and between Korzok and Sakti (p=0.0087). Also, individuals residing in Hanle and Sakti differ significantly in their waist circumferences (p=0.0052) (Table 5b).

**Figure 2:**
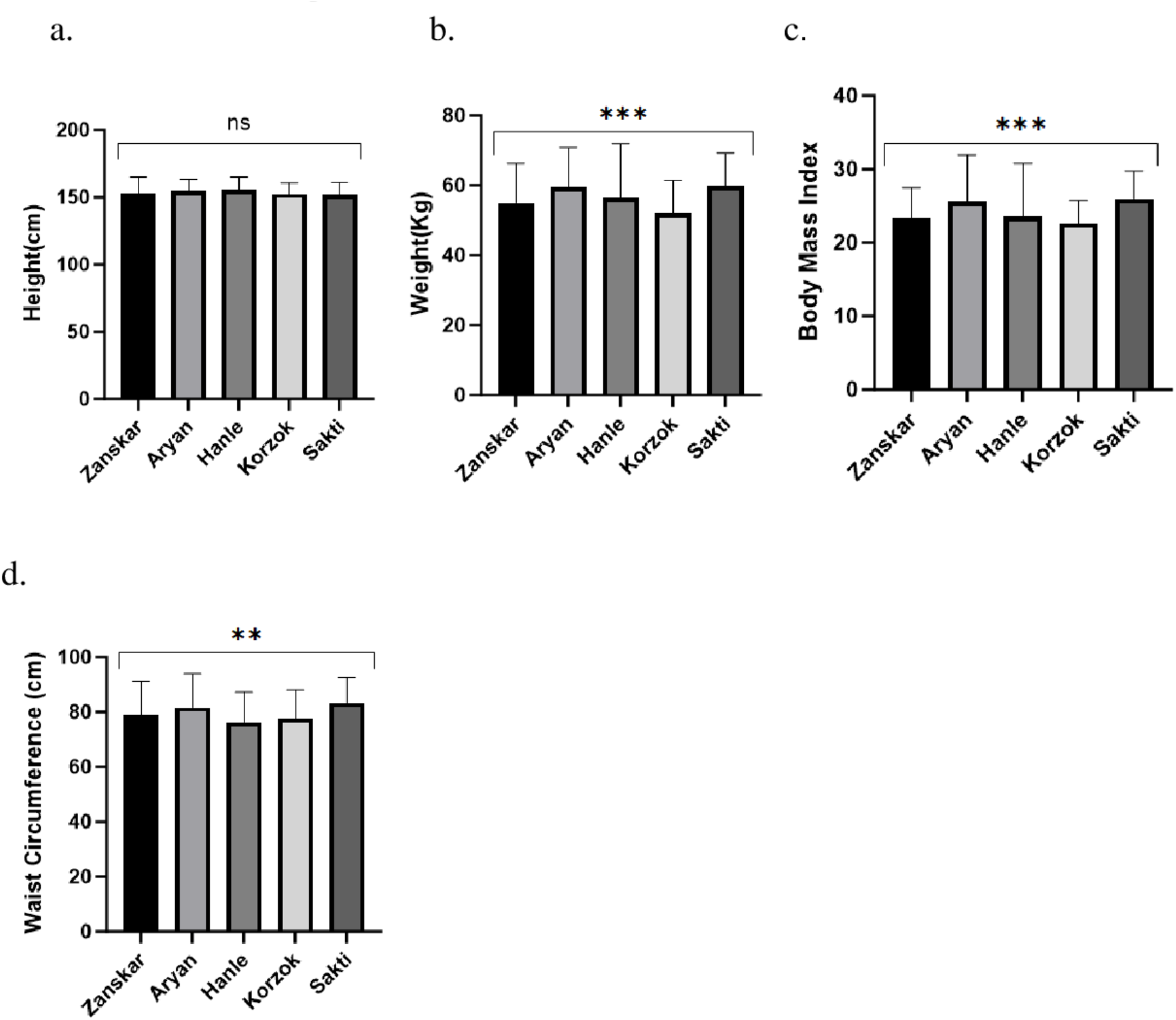
Anthropometric profile study for the distinctive population of ladakh across individuals living in various regions of Ladakh showing a) Height(cm) b) Weight(kg) c) Body mass index d) Waist circumference(cm) (Significance levels ****p <0.0001, *p = 0.01-0.05, ns= non-significant)

### Evaluation of haematological profile

Hemoglobin content varies across different high-altitude populations across the world. In the current study, interestingly, even within the Ladakh region we observed various levels of haemoglobin across the five regions studied. Aryans exhibited lower levels among all populations (13.28±2.13 g/dl), whereas Korzok showed highest hemoglobin content (23.15±29.49 g/dl) thereby attesting to the fact that as we move to higher altitudes hemoglobin increases. Korzok also stands out with higher haemoglobin content both in females and males (17.89±2.98 g/dl and 20.75±2.219 g/dl) (Table 1,2 & 3) when compared to the other groups.

Among Aryan males, the mean partial oxygen pressure is the highest (90.54±2.956%), while Sakti records the lowest level (83.64±6.83%). On considering the mean values, Korzok females (17.89±2.98%) show the highest partial oxygen pressure, while Zanskar females exhibit the lowest level (12.61±2.275%) (Table 1, 2 & 3).An exploration of correlations reveals a non-significant association for partial oxygen pressure. Moreover, a negative correlation (r=-0.4474) (Table 4) is observed between Hanle and Korzok regarding hemoglobin levels, indicating a potential difference between these two regions. The statistical analysis utilizing one-way ANOVA [Figure-3a and 3b)] demonstrates significant results for partial oxygen pressure between Hanle and all other four regions (p<0.001). Similarly, we can see that the individuals across regions vary significantly in their hemoglobin levels (p<0.001) (Table 5b).

**Figure 3:**
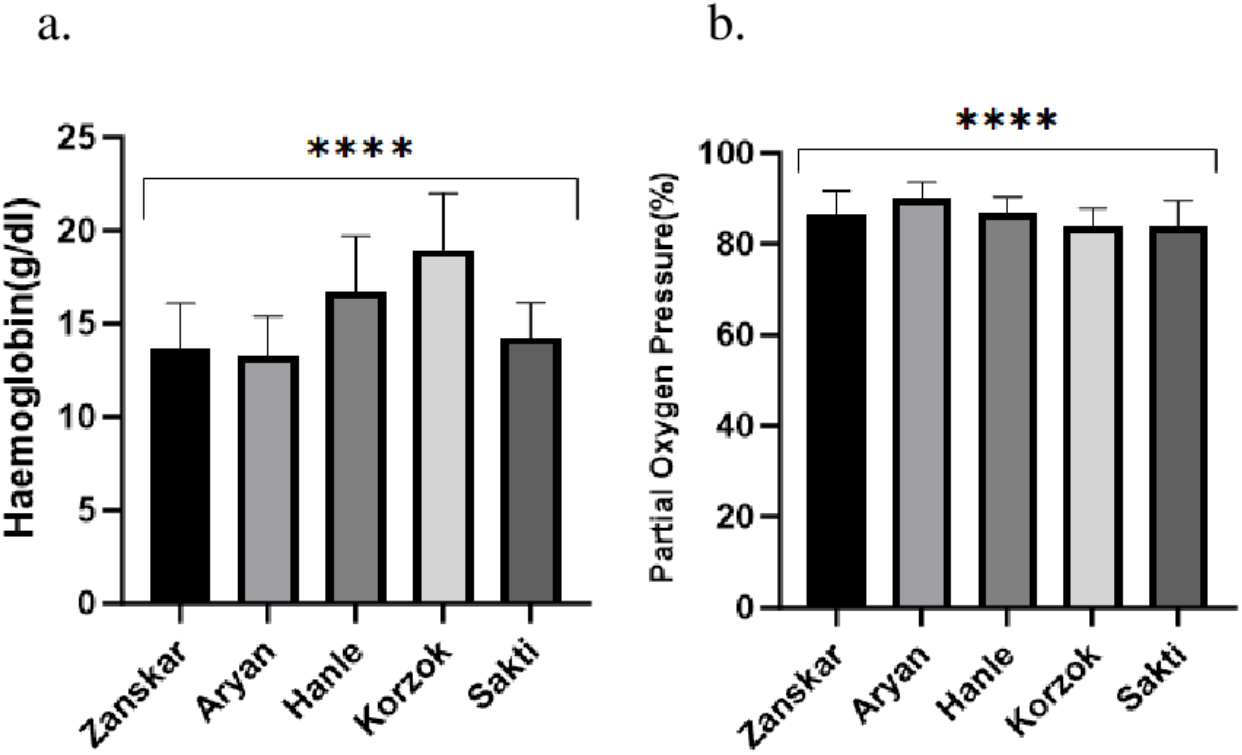
Hematological profile analysis across individuals living in various regions of Ladakh showing a) Hemoglobin (g/dl) b) Partial oxygen pressure(%) (Significance levels ****p <0.0001, *p = 0.01-0.05, ns= non-significant)

### Evaluation of Cardiovascular profile

We find notable variations across regions for systolic blood pressure, with Sakti displaying the highest levels (133.3±25.13 mmHg) and Hanle recording the lowest levels (116.7±20.8 mmHg). In terms of diastolic blood pressure, Aryan stands out with the highest values (85.25±12.2 mmHg), while the lowest levels are observed in Hanle (74.69±13.4 mmHg) (Table 1,2 & 3). When considering gender-specific data, Sakti emerges as the region with the highest systolic blood pressure for males (137.82±15.29 mmHg) and females (131.46±28.042 mmHg). In contrast, Aryan records elevated diastolic blood pressure levels for both males (87.19±13.75 mmHg) and females (84.46±11.58 mmHg)(Table 1,2 & 3). Exploring the correlation analysis, no significant association is identified for systolic blood pressure. However, a negative correlation is observed between diastolic blood pressure in Hanle and Zanskar (r=-0.305) (Table 4).

Utilising ANOVA analysis [Figure 4 (a and b)] with multiple comparisons, statistically significant differences are found. Specifically, for systolic blood pressure, the comparison between Hanle and Sakti yields significant results (p=0.0003). Concerning diastolic blood pressure, significant differences are identified between Zanskar and Hanle (p=0.0009), Aryan and Hanle (p<0.0001), and Hanle and Sakti (p=0.0085) (Table 5b) similar to heart rate difference between Hanle and Korzok showed (p=0.0016) when multiple comparisons were done [Figure 4(c)]. The correlation analysis emphasizes a negative association between diastolic blood pressure in Hanle and Zanskar. Furthermore, ANOVA results underscore significant differences in blood pressure values among various regions, shedding light on potential cardiovascular health disparities. Remarkably, “a positive correlation is identified between Korzok and Zanskar (r=0.3208) (Table 4) in terms of heart rate, and exhibits similar physiological responses of the two regions’’.

**Figure 4:**
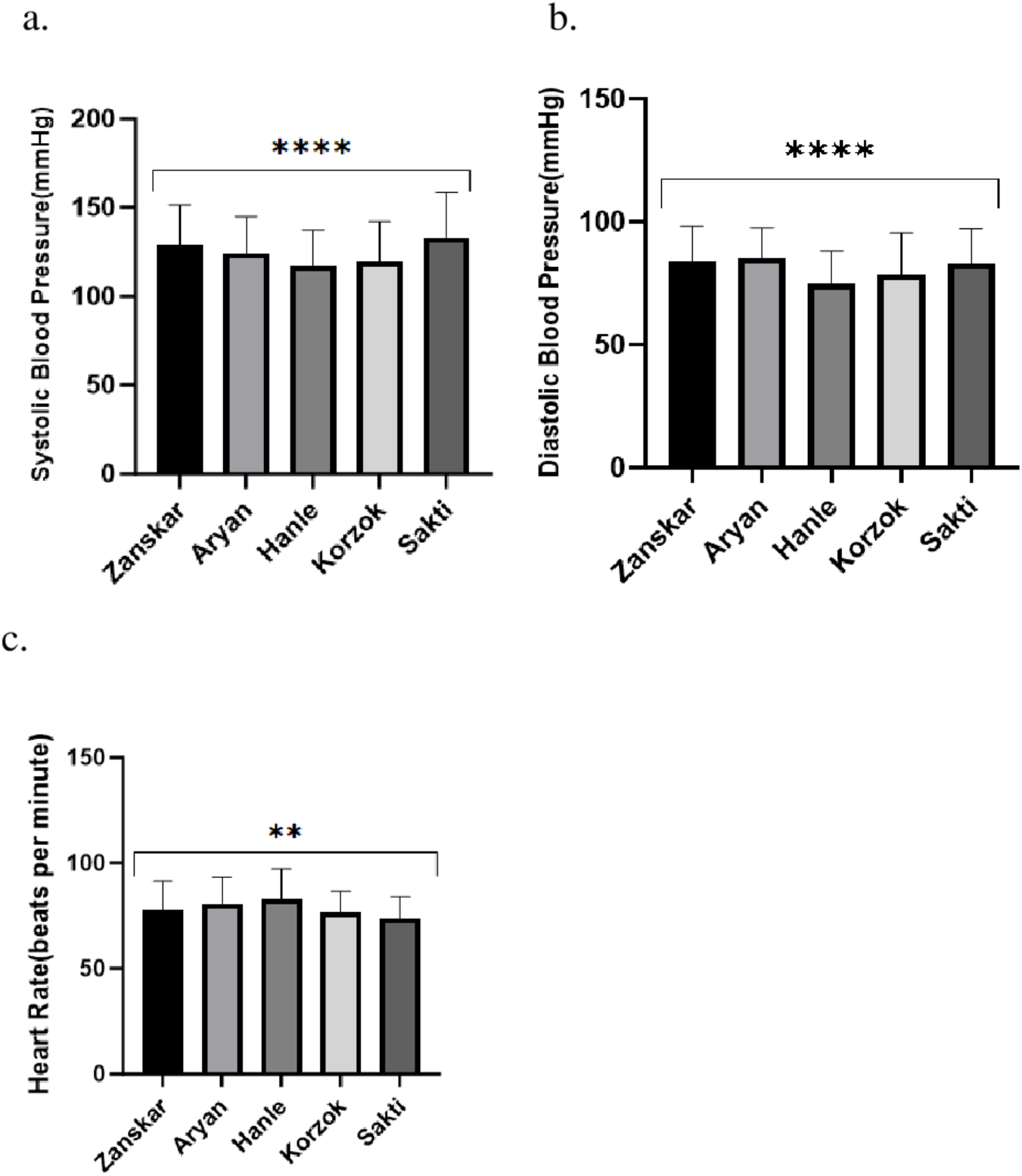
Changes in cardiovascular properties across individuals living in various regions of Ladakh showing a)Systolic Blood Pressure(mmHg) b)Diastolic Blood Pressure (mmHg) c) Heart Rate(beats per minute)(Significance levels ****p <0.0001, *p = 0.01-0.05, ns= non-significant)

### Evaluation of lipid profile

The analysis of cholesterol levels across various regions of Ladakh reveals that Sakti has the highest mean cholesterol value of (200.9±49.95 mg/dl) and the cholesterol values for the normal range are less than 200 mg/dL. This trend is consistent for both males (199.94±55.45 mg/dl) and females (201.25±48.30 mg/dl), with Sakti consistently showing higher cholesterol values than other regions. Interestingly, Sakti also exhibits elevated levels of triglycerides (153.6±67.12 mg/dl) and low-density lipoprotein (LDL) cholesterol (135.8±45.82 mg/dl). At the same time, Zanskar stands out with the highest level of high-density lipoprotein (HDL) cholesterol (43.03±5.483 mg/dl). In comparison, we found that among males, Aryan shows the highest levels of triglycerides (157±54.67 mg/dl) and HDL cholesterol (45.65±3.405 mg/dl). Notably, Sakti demonstrates dominance in LDL cholesterol levels among males (135.23±56.108 mg/dl). Among females, Sakti leads in both LDL cholesterol (136.069±41.82 mg/dl) and triglycerides (155.81±72.28 mg/dl), whereas Aryan exhibits the highest HDL cholesterol level (42.86±2.467 mg/dl). When heart rates were compared, they were highest in Hanle (95.54±105.9bpm) and lowest in Sakti (73.55±10.36bpm). whereas, for heart rate in males(78.86±14.63bpm) and females(103.80±80.59bpm) Hanle showed highest values for both (Table 1,2 & 3).

We then further studied the correlation between regions which revealed non-significant associations for triglyceride and high-density lipoprotein, showed a positive association (r=0.6437) between Hanle and Korzok and for low-density lipoprotein, weakly negative association(r=-0.2562) is seen between Zanskar and sakti valley and for cholesterol weakly negative association(r= −0.2601) is observed between Hanle and Sakti (Table 4). One-way ANOVA analysis showed statistical differences in cholesterol among Zanskar and Aryan (p<0.001), Zanskar and Sakti (p<0.001), Aryan and Hanle (p<0.001) and Hanle and Sakti (p<0.001) and for Triglycerides, similar results were concluded[Figure 5(a and b)]

**Figure 5:**
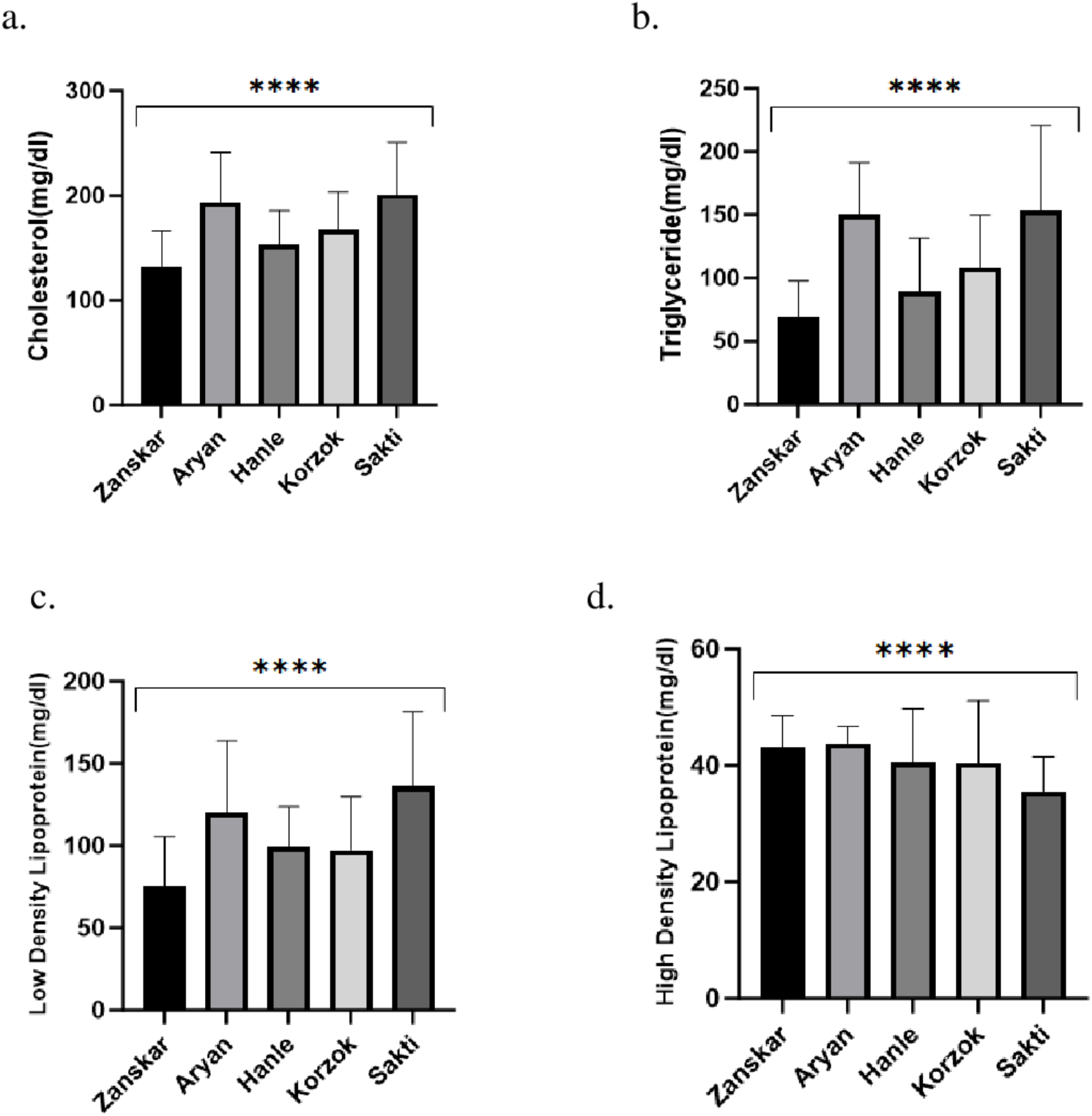
Comprehensive lipid profile assessment across individuals living in various regions of Ladakh showing a) Cholesterol levels b) Triglyceride levels c) Low Density Lipoprotein levels d) High Density Lipoprotein levels (Significance levels ****p <0.0001, *p = 0.01-0.05, ns= non-significant)

In the case of high-density lipoprotein, Zanskar, Aryan and Hanle showed statistically significant values with Sakti (p<0.001). For low-density lipoprotein, Zanskar showed significance when compared with Aryan and Sakti (p<0.001) and Hanle and Sakti showed significant value(p<0.001) (Table 5b) [Figure 5(c and d)].

## 5. DISCUSSION

The analysis demonstrates noteworthy cholesterol and lipoprotein level variations across different regions and genders, with Sakti consistently displaying higher cholesterol, triglycerides, and LDL cholesterol levels. The fact that Sakti has higher cholesterol and triglycerides could also be possibly because its located closer to Leh city and therefore people have more access to processed food. These findings further shed light on potential regional disparities in cardiovascular health markers. Several cases of fatal diseases were also seen in Ladakh highlanders and some parts of North Sikkim due to less consumption of fresh vegetables, fruits and milk and high consumption of smoked meats, which leads to such diseased conditions (De et al., 2023). The Zanskar subdivision, which is located at an intermediate high altitude (3500-3900 m), has a population that is primarily involved in farming and livestock husbandry. The main dietary staples are butter tea, native beverage chang, thukpa, barley flour kholak, rice, and legumes; meat is rarely eaten. Because fresh fruits and vegetables are rarely available in Zanskar and Changthang, hypertension is higher in Sakti (Norboo et al., 2015). . Also, further sedentary occupations add to it. Norboo and his colleagues when compared the city dwellers of Leh and individuals in nomadic area of Changthang found that the former had higher prevalence of hypertension (Norboo et al., 2015). With the fact that higher altitudes generally have lower hypertension than lower altitudes (Tripathy and Gupta, 2007) but in this study this trend is affected due to the lifestyle or close proximity to the city dwellers.

The current study also suggests distinct haemoglobin content variations among different populations. Basak and his colleagues have also earlier reported heterogeneity in the hematological parameters of high and low altitude Tibetan populations (Basak et al. 2021). Here, we also see heterogeneity among the populations studied. Andeans have demonstrated that higher hemoglobin is an adaptation at high altitudes however the same did not hold true for Tibetan and Ethiopian populations (Beall et al. 1998, 2002) . For them the hemoglobin of the individuals living at higher altitude is almost similar to those at the sea level (De et al., 2023). This further suggests difference in erythropoietic response towards altitude exposure in populations which was also reflected in our study. Furthermore, we also observed difference in hemoglobin with increasing altitude in males and females (Supp Fig 1a and 1b). Aryans have lower haemoglobin levels compared to Korzok, with Korzok women having notably higher levels. The average hemoglobin of 17.46 (2.18) g/dl and SpO2 of 84% (9.5%) recorded in Korzok (4550 m) inhabitants is a well-known reaction to low oxygen saturation (Norboo et. al_2010). The analysis of partial oxygen pressure indicates notable differences between regions, with Hanle consistently showing significant variation. Additionally, the correlation between haemoglobin levels further highlights distinctions between Hanle and Korzok. These findings contribute to our understanding of physiological differences across populations and regions. A huge number of separate physiological and biochemical processes work together to keep tissue PO2 levels stable Gender and rising age are related to lower hemoglobin values in Tibetans at high altitude than in Han, and we suggest that genetic variables play a role (Sharma et al., 2022). Lower partial oxygen pressure correlates with higher elevations. Gender and rising age are related with lower hemoglobin values in Tibetans at high altitude than in Han, and we suggest that genetic variables play a role. Furthermore, the EGLN1 and EPAS1 genes exhibit a strong enrichment of high-altitude ancestry in the Tibetan genome, showing that migrants from low altitude acquired adaptive alleles from highlanders (Jeong et al., 2018). In essence, the analysis underscores the prominence of Sakti in terms of body weight, height, and body mass index. The male population of Zanskar displays distinct height and body mass index values, while Sakti’s female contingent exhibits higher measurements in all categories. Moreover, the correlation analysis reveals nuanced relationships, particularly between Korzok and Aryan for height, as well as between Zanskar and Korzok for body mass index. The one-way ANOVA results further validate these distinctions, contributing to our understanding of body composition variations among different groups. In summary, the study highlights distinct patterns in systolic and diastolic blood pressure levels across different regions. Sakti consistently displays higher systolic blood pressure levels, while Aryan shows elevated diastolic blood pressure. People living at high altitudes in cold desert locations have higher diastolic blood pressure and a larger increase in blood pressure with age. rs10495582 is a significant SNP that may be linked to high-altitude essential hypertension (Pandey et al., 2016). Some Highlanders acquire a clinical state known as H-ALT renal syndrome104, which includes intact glomerular function, polycythemia, hyperuricemia, microalbuminuria, and hypertension. Despite the apparent influence of H-ALT on biological processes that govern systemic BP, our understanding of mechanisms leading to or changing the prevalence and clinical course of chronic hypertension at H-ALT is inadequate (Narvaez-Guerra et al., 2018). Basically, the analysis underscores the diversity in heart rate across regions, with Hanle showcasing elevated levels. Gender-wise, Hanle maintains its lead in heart rate. It can be concluded that there are some factors involved in order to cause cardiovascular risks in the populations residing at particular altitudes. The study emphasizes the dissimilarities between Hanle and Sakti for waist circumference, offering insights into the physiological differences in these regions. In examining body weight variations among the high-altitude populations of Ladakh, significant associations were explicitly observed between individuals from the Aryan and Korzok areas. This finding suggests that geographical and possibly genetic factors contribute to differences in body weight among populations in these distinct regions. Adapting to high-altitude living may differently influence metabolic and physiological traits, including body weight, across these communities.

### Limitations

We would like to point out that we could not complete the genotyping studies due to a lack of funding. However, we will undoubtedly continue the work if appropriate funding becomes available. In addition, we have data from only five regions of Ladakh.

### Conclusion

There are various reasons why the morphological and physiological domains alter when we move from lower to higher elevations, whether they are local or throughout the region. One of these elements could be the availability of food resources, which impacts their dietary mechanisms and contributes to other aspects that further assist with differences in lifestyle. Furthermore, a profound phenotypic analysis reveals gender differences in population sizes across all locations and heterogeneity in physiological responses towards high altitude.

## Acknowledgement

The authors would like to thank all the volunteers who participated in the study. We would also like to acknowledge Ladakh Institute of Prevention, Leh for their support in sampling collection. C.B.M was supported by DBT/Wellcome Trust India Alliance Early Career Fellowship (IA/E/18/1/504338). GC is supported by ICMR ad hoc grants ICMR ad-hoc grants (2021-6389), (2021-11289) and BHU IoE incentive grant BHU (6031). We also thank our fellow researchers for their valuable insights, discussions and critical comments, and we would like to express our gratitude to LIP Staff members Mr Mohammad Iqbal, Mrs Sherab Dolma, Mrs Tsering Spalzes, Mrs Jigmet Ladol and Tsering Dolker.

## Author contributions

BC, UC, CBM and GC have worked on the study design and conceptualized the study. UC, NPW and TN has collected the data, done the field work, provided inputs about the populations in the study and assisted in writing the manuscript. BC has done the phenotype assessment, data analysis and interpretation and drafted the original manuscript. TN and NPW has helped in data curation and field work. GC has reviewed and edited the manuscript. CBM has supervised the study, worked on the methods, and reviewed and edited the manuscript.

