## Supplementary figures and images for "A sneak peek at the high altitude adaptation of the Ladakh populations"

### Supplementary figures (1).tiff

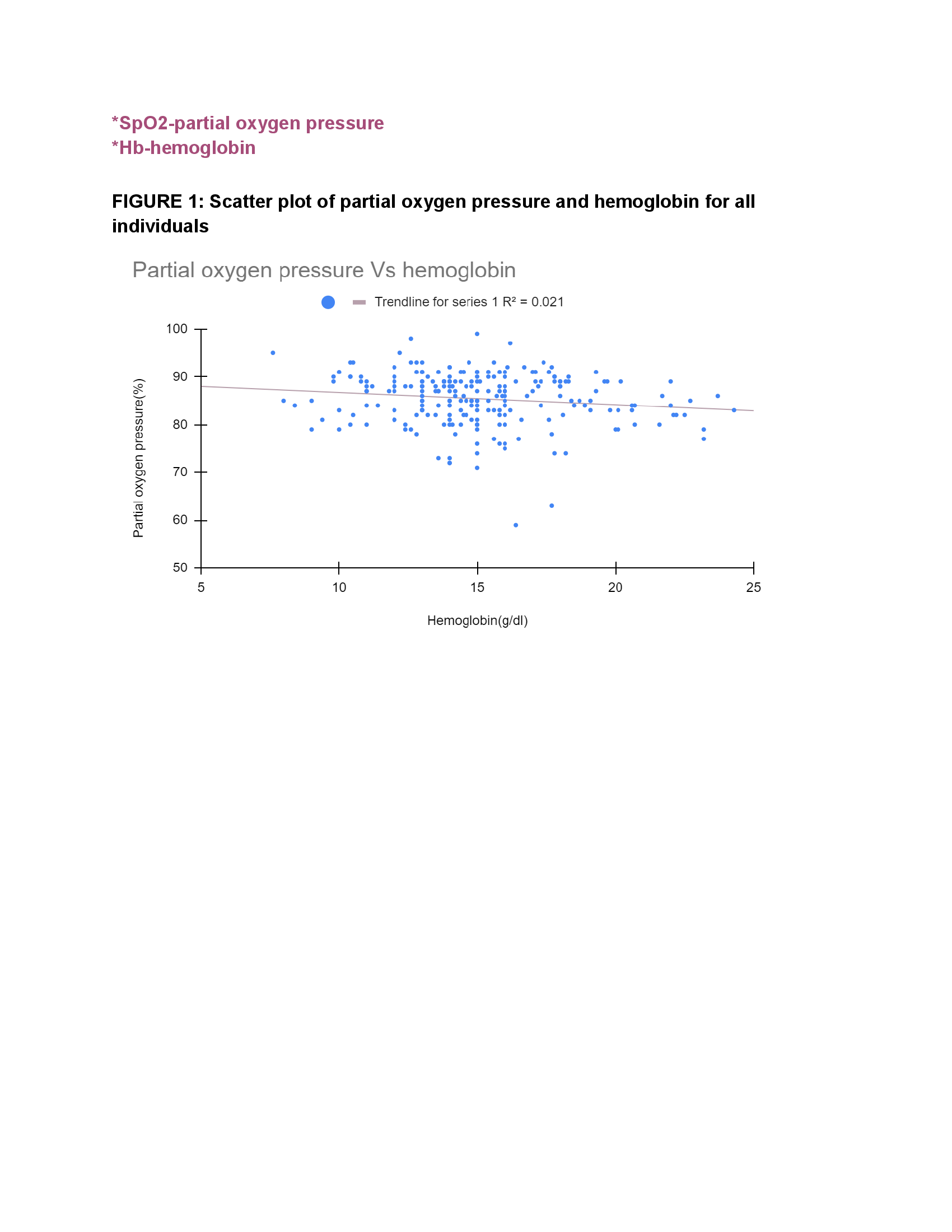
